# Sublethal Interaction Factor (SIF), a growth-based method to analyse antibiotic combinations at sub-inhibitory concentrations

**DOI:** 10.1101/2025.03.04.641412

**Authors:** D.L. Sánchez-Hevia, N. Fatsis-Kavalopoulos, D.I. Andersson

**Affiliations:** Department of Medical Biochemistry and Microbiology, Uppsala University

## Abstract

Antibiotic resistance is a global concern with significant implications for healthcare, food production and the environment. Therapies involving antibiotic combinations are frequently employed as a strategy to overcome antibiotic resistance. When present in combination, the efficacy of antibiotics may be enhanced or weakened, and as antibiotic interactions are *a priori* generally unpredictable they need to be experimentally determined. Though antibiotics are regularly present at sublethal concentrations (for example in patients with suboptimal dosing regimens and in the environment), effects of antibiotic combinations are generally studied at lethal dosages. To address this, we developed the Sublethal Interaction Factor (SIF) assay, based on the Bliss Independence model, to quantify antibiotic combination effects at sublethal concentrations. SIF assay uses the whole growth curve, instead of only the growth rate, and determines reliably the outcome of the interaction between two antibiotics at sub-lethal concentrations. The SIF method was validated against checkerboards and CombiANT assays and showed high sensitivity and specificity, attesting to its usability.

## Introduction

Antibiotic resistance is a global threat, jeopardizing the efficacy of antibiotics in treating bacterial infections. This, along with the existing antibiotic discovery void, has made relevant the use of treatment with antibiotics in combination. Antibiotics have also been detected in a wide range of environments^1–7^, with several natural habitats identified as antibiotic resistance reservoirs^7–9^. In all those cases antibiotics are not present individually, but as combinations (and for the environment also other pollutants). The efficacy of those antibiotic mixtures may range from an effect equivalent to the sum of their individual activities (additivity), to an enhanced effect (synergy) or a reduced effect (antagonism)^10–12^. Even though the use of antibiotic combinations is not new, some date back to the 1950s^13^, in many cases little is known regarding the mechanisms underpinning the observed interactions. Many efforts have been made to predict antibiotic interaction effects^14–16^; but the species- and strain-dependent nature of antibiotic interaction outcomes^17–19^, along with the demonstrated impact of genetic factors on drug interactions^20^, underscores the continued relevance of in vitro analysis.

To date, there is no consensus on the best procedure to quantify an antibiotic interaction. Different laboratories not only use different experimental techniques, but also different models to analyse the data, such as the Loewe and Bliss models^16,21–25^. The model proposed by Loewe posits that an antibiotic cannot interact with itself, whereas the Bliss independence model assumes that the effect of a drug combination is the product of the single-drug effects. Both Loewe’s and Bliss’ models have limitations^16,22,25^, and Loewe’s model is the preferred one when analysing two antibiotics which share a target^20,26^, whereas the Bliss model is used when two compounds act on different targets^12,16–18,27,28^.

Several established assays, such as time-kill, checkerboard^26–31^ and CombiANT^19,34–36^ assays, are commonly used to study antibiotic interactions, all of which are based on lethal antibiotic concentrations. Both time-kill and checkerboard assays can quantify the overall growth of the cultures, but the standardized determination is typically done as an endpoint comparison. For time-kill assays an increase or decrease as >2 log^10^ CFU/mL from the reduction achieved with most active agent is set to determine if an interaction is antagonistic or synergistic, respectively^33,37^. In the case of checkerboard and CombiANT assays, the quantification is done by the Fractional Inhibitory Concentration Index (FICi; formula 1)^32–34,38^; considering the MIC values of each antibiotic tested alone, as well as their Inhibitory Concentration (IC) tested in combination.

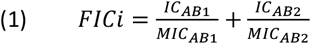

Since a drug cannot interact with itself, a self-drug combination will always be independent (additive) with a FICi=1, and deviations from additivity entail interactions, distinctly FICi<1 and FICi>1 for positive and negative interactions respectively^19,38^. However, in practice, only interactions with a FICi≤0,5 or FICi≥4,0 are regarded as synergistic or antagonistic^19,32–34^.

Despite the widespread use of FICi-based methods, its limitation as a single-point measurement remains a significant drawback since it fails to account for physiological influences during early growth stages, such as the lag phase. A more comprehensive approach to capture interaction effects across all growth phases is to calculate the Area Under the Curve (AUC) for different conditions and compare them, providing a more complete assessment of antibiotic interactions. In this sense, some recent works have used the AUC of cfu (AUCFU) as an alternative method to assess the total bacterial burden over the duration of the study ^25,39,40^. However, no definitions for synergy, additivity and antagonism have been defined for AUCFU^25,39^.

There exist several motivations to determine interactions at sub-MIC. First, it has been shown that antibiotic resistance can be selected at very low concentrations^4,41–45^, and when those interactions are synergistic they may in fact promote antibiotic resistance evolution^28,46^. Second, it has also been shown that the nature of the drug interaction can often be affected by dosage^47^. For sub-MIC assessment, today the preferred approach is Bliss Independence Method (S), which compares the exponential growth rate (g) in presence of agents (here AB_1_ and AB_2_) with the control (here noAB; formula 2)^27,32^. However, as previously noted for FICi, the Bliss Independence Model is limited by its reliance on a single growth phase - specifically, the exponential phase - potentially overlooking critical physiological effects that occur throughout bacterial growth.

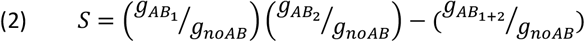

Here we present the SIF (Sublethal Interaction Factor) assay, an easy approach for evaluating antibiotic interactions at sublethal concentrations. Unlike the widely used Bliss Independence method^12,17,27^, which relies on exponential growth rate measurements, SIF offers a distinct advantage by providing an assessment of interactions at sublethal levels based on the whole growth curve. This novel approach enables a more nuanced understanding of antibiotics interactions beyond conventional methodologies.

## Results

SIF is a comparative metric based on the relative growth of bacterial cultures that are exposed to subinhibitory antibiotic concentrations at four conditions: sub-inhibitory concentration of each antibiotic separately, a sub-inhibitory concentration of the antibiotic combination and a control without antibiotics (from now referred as noAB; Fig 1.1). SIF is easily performed in microtiter plates where the four different conditions are established in individual wells. The antibiotic concentration used depends on the strain, the antibiotic and the antibiotic interaction, and must be optimized for each case. Each antibiotic must be at a concentration high enough to produce a measurable effect on growth rate, yet not so high that the culture is eradicated when exposed to both antibiotics in combination. Based on our experiments, the best results come from the use of 0.25x to 0.5x MIC dosages.

**Figure 1.**
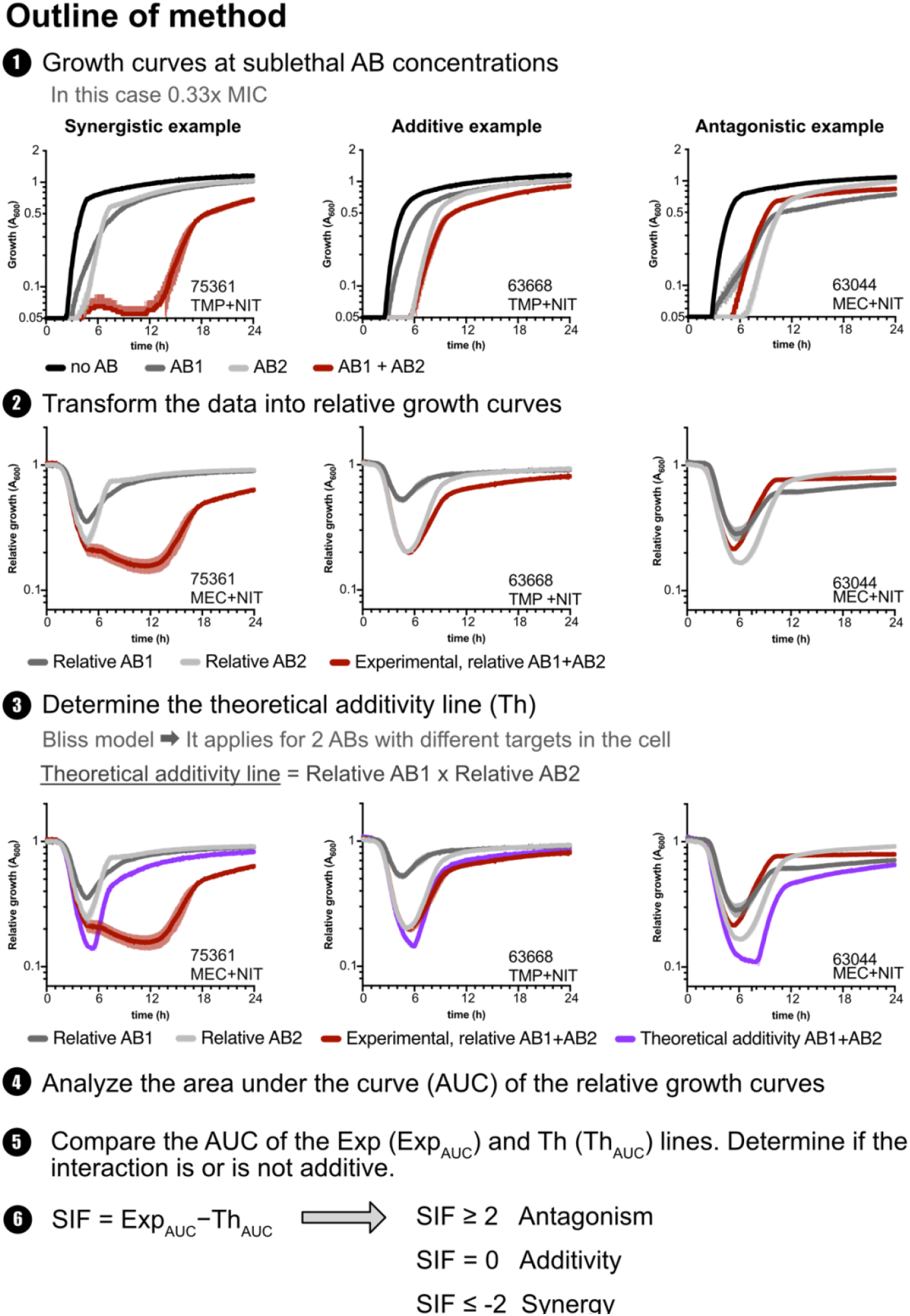
Brief explanation of determination of antibiotic interaction by SIF. Examples for the three possible outcomes (synergy, additivity and antagonism) are shown.

SIF analysis starts by transforming the complete growth curves into relative growth curves by normalizing them with the growth in absence drugs (Formula 3). At this point visual differences between the three different outcomes – synergy, additivity and antagonism – can already be observed (Fig 1.2).

**Figure 2.**
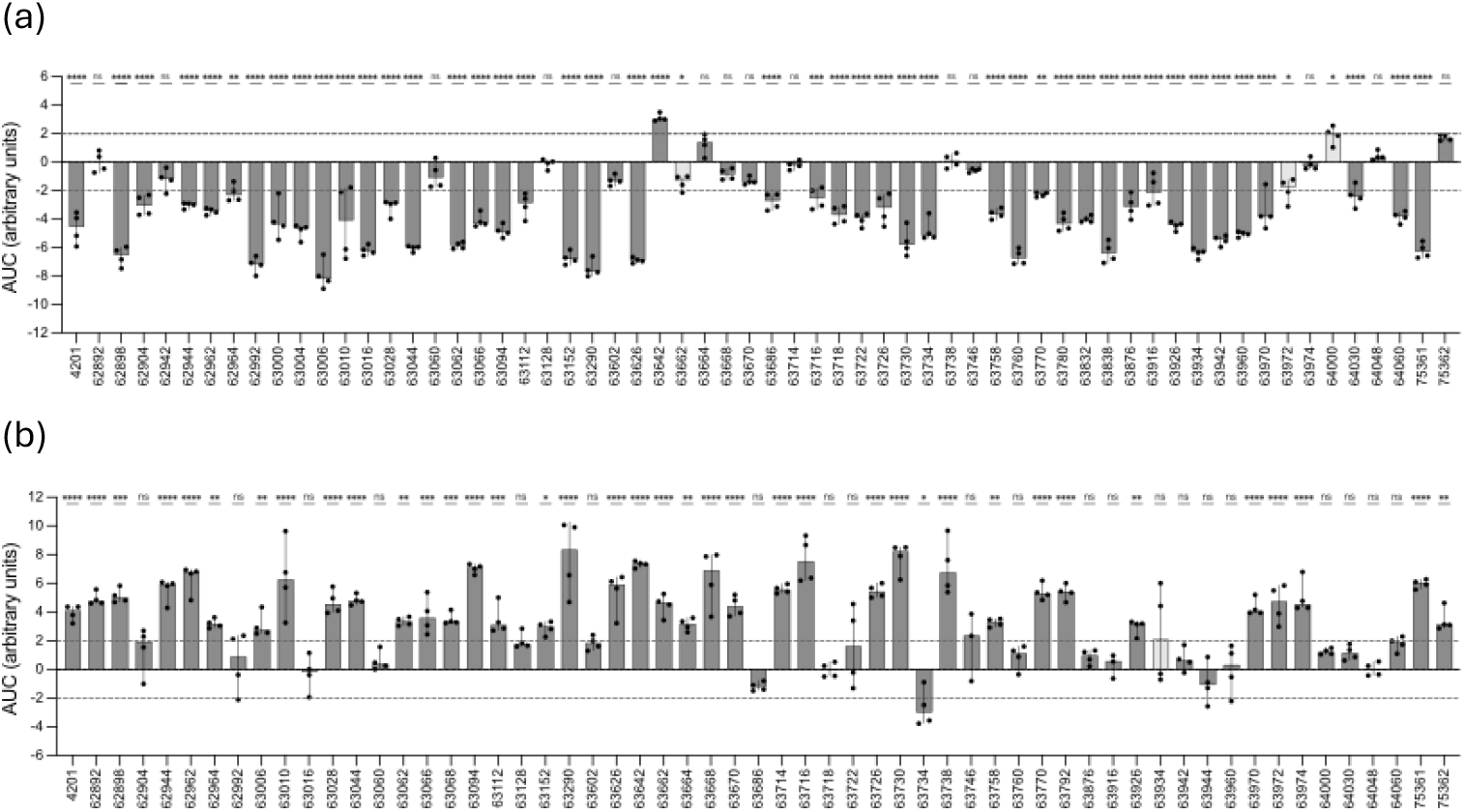
Plot of the SIF result for each of the isolates tested for (a) TMP+NIT interaction TMP+NIT, (b) and for MEC+NIT interaction. The median and the deviation correspond to four independent biological replicates, showed with dots. The statistics indicate if significant differences were found for each case when experimental (Exp_AUC_) and theoretical (Th_AUC_) area under the curve values were compared. The four cases categorized differently by SIF and the statistics are indicated in light grey.

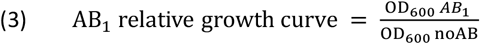

To determine the type of interaction, one compares the relative growth curve in the presence of both antibiotics (from here on referred to as experimental growth or Exp) with the theoretical additivity line or Th (Fig 1.3). Th corresponds to the relative growth rate expected at each time point, assuming a Bliss independence interaction^21^ (Formula 4, Formula 6). If two antibiotics have a positive interaction, the inhibitory effect is enhanced and the Exp line will be below the Th. In contrast, when two drugs show a negative interaction the Exp line is above the Th line (Fig 1.3). Finally, in an additive interaction the combined inhibitory effect is equal to the sum of their individual inhibitory effects, and Exp and Th lines overlap or are close (Fig 1.3).

**Figure 3.**
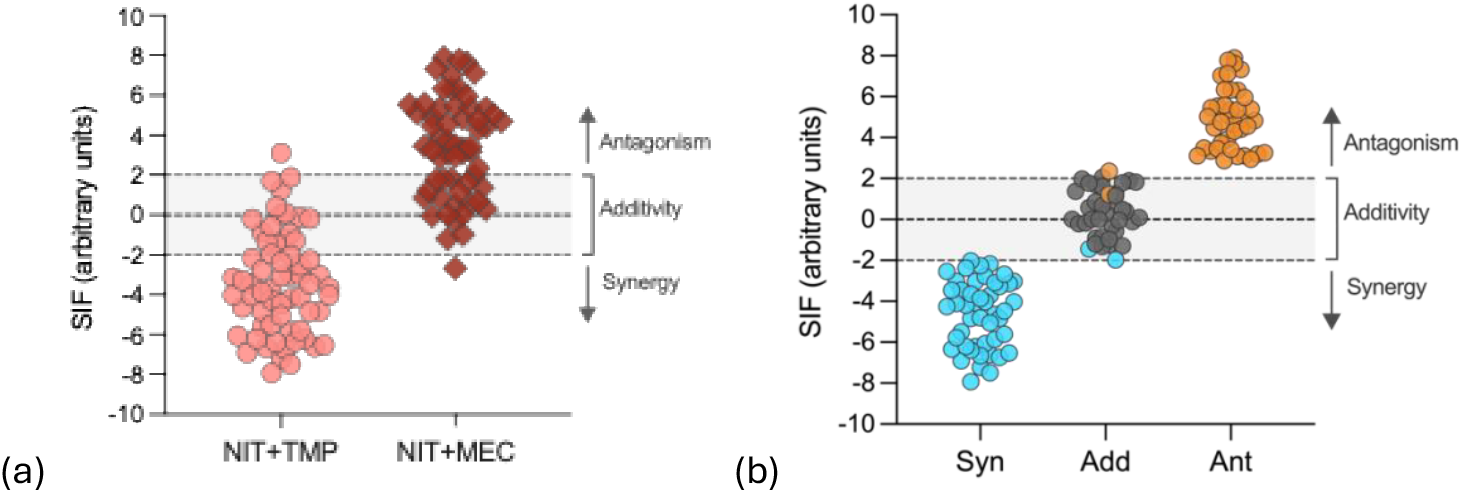
Plot of the SIF result of all the isolates tested, (a) according to the antibiotic combination, or (b) antagonistic (in orange, ANT), additive (in grey, ADD) or synergistic (in blue, SYNG) according to the SIF value and the significant differences. The colour is based the SIF result. Only the isolates which showed a SIF value larger than 2-fold and significant differences between Exp_AUC_ and Th_AUC_ were considered synergistic or antagonistic.

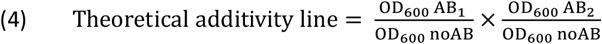

If one compares two conditions where clearances is observed, as it happens during lag phases, the turbidity difference can be equal to 0 or even negative, due to the typical 2% error of the plate readers (suppliers’ information). We propose including a detection limit, such 0.05 in this example (Formula 5), to avoid negative values which would interfere with the following calculations. This matter has more importance in the Th calculation, where a minimal value is of the outmost relevance. In a hypothetic condition and stage where growth is only detected in plain broth - this is, no growth detected in any drug condition neither individual nor in combination -, Formula 4 assumes a repression in drug combination below clearance, which is not physiologically possible. To address this issue, we include the limit of 0.05 in the Th calculation (Formula 6).

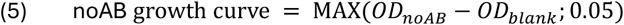

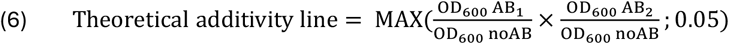

A mere visual comparison of Exp and Th does not allow a precise quantification of the interaction and to overcome this issue one calculates the AUC of the Th and Exp curves (by integrating the respective two plots), thereby converting time curves into a single numerical measurement (Fig 1.4). Subsequently, the AUC of the Exp and the AUC of the Th curves are compared using nonparametric paired t-tests (Fig 1.5). If the two measurements differ significantly (p< 0.05) then one can conclude that the antibiotics interact, and that the interaction is non-independent. The outcome of the antibiotic interaction can be determined by erasing Th_AUC_ from Exp_AUC_ (Formula 7).

**Figure 4.**
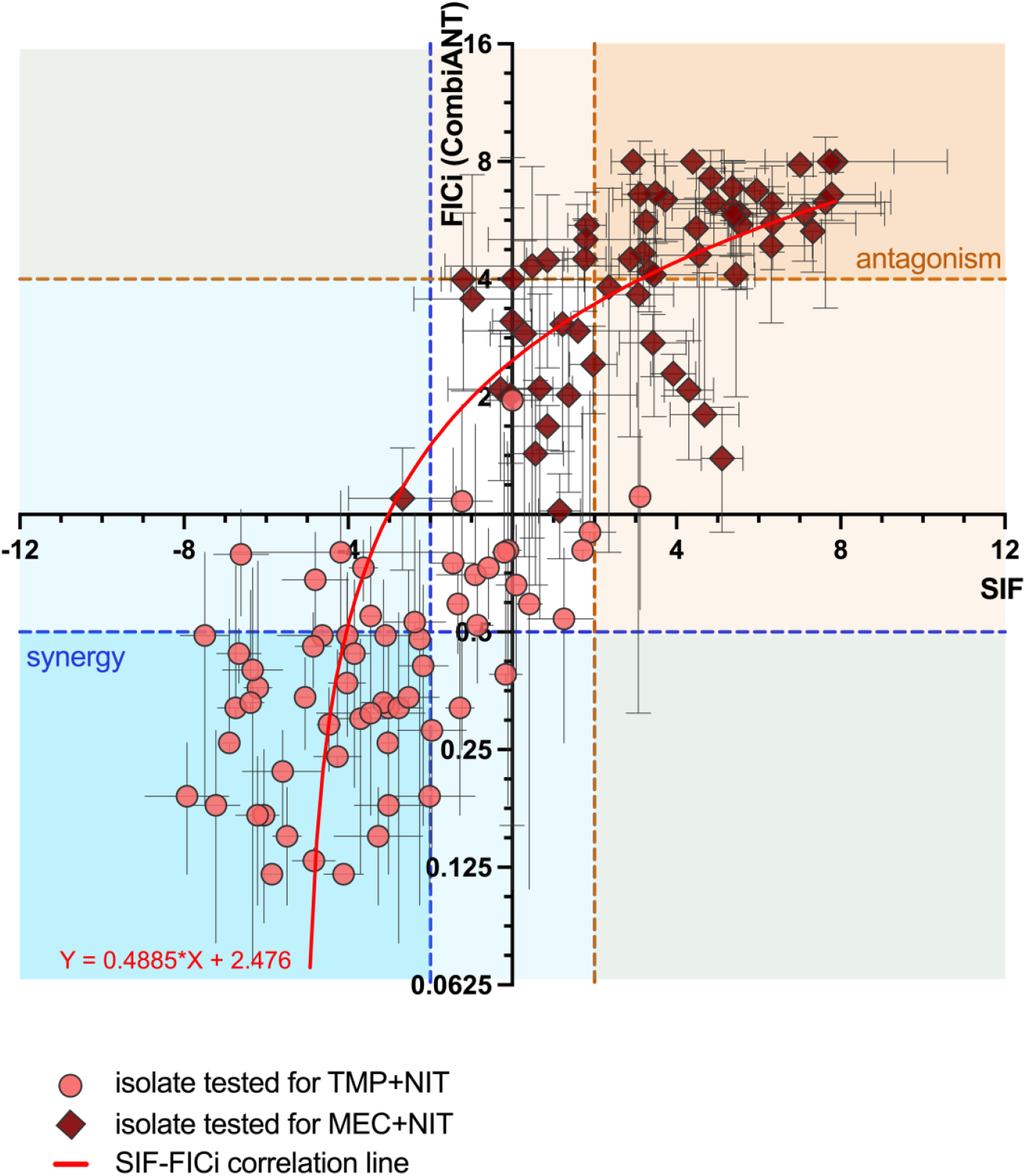
Correlation between SIF and FICi results, showing both the average and the standard deviation values of each isolate. The FICi results were analysed by CombiANT with at least three biological replicates. Both TMP+NIT and MEC+NIT interactions are represented, with circles and diamonds, respectively. The correlation (p<0,0001) was verified by a two-tailored correlation test.

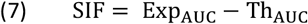

To verify that our data was normally distributed and could by analysed by a nonparametric paired t-test, we performed a normality test using 12 independent assays of the DA75361 strain exposed to the NIT+TMP combination. Both the AUC values of each experimental condition, the additivity theoretical line and the SIF values passed the tests performed (Table S2, Fig S1).

Considering that for independence the antibiotics do not interact, additivity is when SIF=0, whereas SIF>0 and SIF<0 correspond to negative and positive interactions, respectively (Fig 1.6). Since not all positive or negative interactions necessarily qualify as synergistic or antagonistic, one needs a clear and reliable threshold to define these interactions. To establish such a threshold, we analysed the statistically differences in Exp_AUC_ and Th_AUC_ within our dataset. We noticed that interactions classified as synergistic or antagonistic^12,34^ consistently deviated from SIF = 0 by at least a 2-fold change (Fig 2). Based on this finding, we recommend using SIF ≥ 2 to define antagonistic interactions and SIF ≤ -2 to define synergistic interactions, ensuring a robust classification framework.

The SIF assay was tested by determining the antibiotic interactions of trimethoprim (TMP) plus nitrofurantoin (NIT), and mecillinam (MEC) plus NIT in a group of clinical *Escherichia coli* isolates. We chose these drugs for having divergent modes of action and for having been previously described as mainly synergistic and antagonistic, respectively, based on checkerboard and CombiANT assays^12,34^. A total of 121 cases were tested, 62 strains for the combination of TMP+NIT and 59 for the combination of MEC+NIT. Growth curves were obtained at several sub-inhibitory concentrations and we selected 0.33x MIC as a favourable concentration to observe a clear effect on growth when each antibiotic acted individually, but not complete growth inhibition when present in mixture. Each case was assessed at the four conditions: without antibiotic (noAB), only AB1 (TMP or MEC) at 0.33x MIC, AB2 (NIT) at 0.33x and the AB1+AB2 in combination. All analyses and calculations were done as described in Fig 1. Overall, we found smaller Exp_AUC_ for TMP+NIT (Fig. S3a) and larger Exp_AUC_ for MEC+NIT (Fig S3b), indicating their synergistic and antagonistic nature, respectively. Those results were confirmed by SIF determination (Fig 3a) where 68.6% of the isolates showed antibiotic interactions (either synergy or antagonism) and the remaining 28.1% showed additivity, in agreement with the previous work (Fig S2).

To assess the accuracy of the SIF results we compared them with FICi. We decided to employ CombiANT, a previously described and validated agar plate-based method of quantifying interactions^34^, because of its ease of use. We successfully tested 120 out the 121 original cases. Our results demonstrate that the two assays consistently produced similar outcomes (Fig 4), with 80.83% of the cases classified identically by both SIF and FICi (Fig S4). Notably there were no instances where the two methods assigned conflicting classifications (i.e., one categorizing an interaction as synergistic while the other labelled it as antagonistic). This level of correlation between SIF and CombiANT is equivalent or higher than the correlation previously observed for time-kill and checkerboard^29,48,49^, the most commonly used techniques; supporting SIF as a reliable method to explore drug combinations at sublethal doses.

SIF evaluates each case by two criteria, a statistically significant difference between Exp_AUC_ and Th_AUC_ to classify a case as an interaction, and secondly, by a ≥2-fold change in SIF value, to define the type of the interaction. The majority of the cases tested here fulfilled both criteria (96.7%), with 4 of them (3.3%) only fulfilling the first but not the second criterion (Table S1, Fig S2). A closer look of these discrepancies reveals that all of them correspond to cases with a SIF was smaller than 2-fold and whose variation in AUC was found statistically significant (Fig 3b). Such results are to be interpreted as an additive interaction with a significant statistical confidence. When compared to the FICi results for the same strains, specificity was found to be 90.6% and sensitivity 85.0 % (Fig S5).

## Discussion

Understanding the efficacy and selection dynamics of antibiotic combinations under conditions where complete bacterial growth inhibition is not achieved is important not only in the environment, where low antibiotic concentrations are well-documented^1,3,4^, but also in clinical settings. During treatment, antibiotic levels in the body can drop below the MIC due to factors such as suboptimal dosing regimens, patient disease state or poor patient compliance, underscoring the importance of studying sublethal antibiotic interactions^50–53^. To facilitate measurements of antibiotic interactions at subinhibitory concentrations we developed the Sublethal Interaction Factor (SIF) method.

SIF classified AB interactions consistently equivalent to FICi, proving to be a reliable alternative to estimate drug interactions. Moreover, SIF is strict in the assessment of an interaction as non-independence, and only those pairings that exhibit a statistically significant difference between Exp_AUC_ and Th_AUC_, and have a SIF value exceeding the defined threshold, are classified as true interactions (i.e. non-additive or independent), being either synergistic or antagonistic (Table S1, Fig 3b). The high accuracy of SIF (80.8%; Fig S5), is reflected in both the true-positive rate (or sensitivity; 85.0%), and true-negative rate (specificity; 90.6 %).

SIF determines antibiotic interactions at concentrations where only partial growth inhibition occurs, integrating the effect over all stages of growth in the AUC analysis. Other traditional methods based on the estimation of the exponential growth rates may fail in detecting the effects on the lag phase or the final yield; and the analysis becomes complicated when a bacterial strain shows a non-sigmoidal growth curve in presence of antibiotic, as observed for some β-lactams which show pre-lytic OD increase^54^. SIF overcomes this limitation by evaluating the effect of drug combinations over the whole growth curve, achieving a more complete assessment of the interactions. As demonstrated here with the example of mecillinam (MEC), SIF is equally suitable for antibiotics that affect the exponential phase or disrupt the normal sigmoidal growth pattern. Additionally, unlike previous analyses based on AUCFU^25,39,40^, SIF integrates growth over the entire curve while incorporating well-defined boundaries for different antibiotic combination outcomes. Consequently, SIF provides a refined reinterpretation of previous approaches to assessing drug combinations, unifying the analysis power of the AUCFU methods with a more precise characterization of combination outcomes.

Important advances have been made in understanding the impact of low, or very low, antibiotic concentrations on resistance evolution, observing Minimal Selection Concentration (MSC) as low as 1/2000 of MIC for some antibiotics^41–43,46^. However, the impact of this low concentration in drug combinations remains unclear. What is more, the study of drug interactions at sub-inhibitory concentrations is also crucial from both ecological and evolutionary perspectives. Even if synergy at sublethal concentrations does not completely inhibit bacterial growth, it can still accelerate the rate of evolution of resistance^28,46^. Therefore, the influence of diverse antibiotics, as well as other pollutants, such metals and antibacterial biocides, on bacterial populations in natural habitats remains an active area of research^55,56^. Furthermore, it has been shown that antibiotic susceptibility can be influenced by changes in carbon sources and availability of micronutrients (i.e. iron)^57–59^-as reflected in the higher production of ß-lactamases in rich medium as compared to minimal medium^60^ -, or by the activation of stress responses^45,60–62^. However, how these conditions alter the antibiotic interactions is not well explored and SIF can provide a robust and adaptable method to test different growth conditions.

In this study, we have presented a method that fills a gap in our arsenal of laboratory assays for determining the effects of drug combinations. SIF is highly accessible, its implementation requires only standard laboratory equipment, and its analytical process is straightforward yet robust, making it adaptable for microbiology laboratories of all scales. Unlike conventional methods that rely on single-point measurements, SIF captures interaction dynamics across the entire bacterial growth curve, providing a more comprehensive assessment of antibiotic effects.

Finally, beyond its immediate applications, SIF represents a step forward in refining how we analyse drug interactions in both clinical and environmental contexts. Its versatility extends beyond antibiotics, offering a framework for studying the combined effects of other antimicrobial agents, such as heavy metals, biocides, and antimicrobial peptides.

## Material and Methods

### Bacterial strains and culture media

The *E. coli* strains used in this work are K-12 MG1665 (referred to here as 4201) and bloodstream infection isolates^63^. Bacteria were cultivated either in Mueller-Hinton broth (MHB; Becton Dickinson) or on Mueller-Hinton agar (MHA; Becton Dickinson) and supplemented with antibiotics when indicated. The antibiotics used in the assay are: Mecillinam (MEC; Sigma), Trimethoprim (TMP; Sigma) and Nitrofurantoin (NIT; Sigma). The antibiotics stocks were prepared using Dimethyl sulfoxide (DMSO, Sigma-Aldrich) or deionized water (Sigma-Aldrich) following supplier’s recommendations. Bacterial cultures were incubated at 37 °C and growth was monitored by measuring turbidity at 600 nm (OD_600_). MIC determination was done with E-test strips (Biomerieux).

### Antibiotics interaction determination by CombiANT

FICi and the interaction profile was quantified by CombiANT (Rx Dynamics AB, Uppsala, Sweden), following the previously described protocol^19,34^. At least three independent biological replicates were tested, the FICi was calculated by formula 1, and the average of all the replicates was determined.

### Sublethal interaction factor (SIF)

The SIF assay compares the growth of the cultures at four conditions: broth (noAB), medium with one antibiotic (either AB1 or AB2) or the two antibiotics in combination (AB_1_+AB_2_). Antibiotics are supplemented at sub-inhibitory concentrations, typically between 0.25x to 0.5x MIC dosages. Here, MEC, TMP and NIT were supplemented at 0.33x MIC. Each isolate was tested with a minimum of four independent biological replicates. The overnight cultures were diluted in PBS (phosphate-buffered saline buffer), and 10_5_ cfus were inoculated in each well. The assay was performed in honeycomb Bioscreen plates (Bioscreen Inc.). After inoculation, the plates were incubated at 37 °C in the Bioscreen (Bioscreen C; FP-1100-C), and the OD_600_ was monitored every 4 minutes for 24 hours.

The analysis of the interaction was done as follows (Fig 1). First, the growth curves were transformed into relative growth curves, by dividing the turbidity in the presence of AB_1_ and/or AB_2_ to the growth of the culture at plain media (Formula 3). Then, following the Bliss model_21_, the Theoretical additivity line (Th) was determined (Formula 4), which will work as antibiotic interaction threshold.

One further calculates the AUC of the experimental relative growth (Exp_AUC_) and the theoretical additivity line (Th_AUC_). The analysis of the Area Under the Curve (AUC) was done in lineal scale, and a trapezoid integration method was used. Time-point 0 was excluded as relative survival rates do not apply in the beginning of the experiment. Subsequently, the paired t-test was applied to determine differences between Exp_AUC_ and Th_AUC_. Cases deemed statistically significant are associated with deviations from independence; that is, AB1 and AB2 interact, irrespective of whether the interaction is positive or negative. To further assess the type of interaction, one determines the SIF value (Formula 7).

### Statistical analyses

Statistical analyses were performed using Graph Pad Prism; ns indicates non-significant, * p<0.03, ** p<0.002, *** p<0.0002, *** p<0.0001 for all hypothesis tested in this work.

## Declaration of interests

We declare no competing interests.

## Funding

This research was funded by grants to DIA from Swedish Research Council 733 (grant nr 2021-02091), Wallenberg Foundation (grant nr 2018.0168) and 734 Swedish Foundation for Strategic Research (grant nr ARC19-0016).

## Author contributions

Conceptualization: DLSH, NFK, DIA

Investigation: DLSH, NFK

Methodology: DLSH Visualization: DLSH

Software: DLSH, NFK

Writing original draft: DLSH

Writing review and editing: DLSH, NFK, DIA

Funding acquisition: DIA

## Notes

### Competing Interest Statement

The authors have declared no competing interest.

