## Supplemental figures for "Sublethal Interaction Factor (SIF), a growth-based method to analyse antibiotic combinations at sub-inhibitory concentrations"

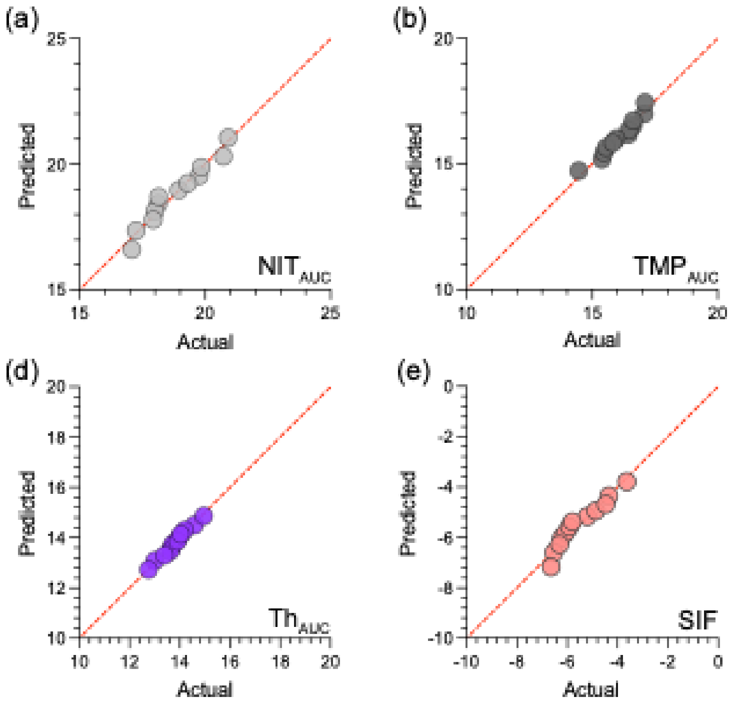


Supplementary figure S1: Normal Q-Q plot of the 5 conditions analysed by the normalized test (a) $NIT_{AUC}$, (b) $TMP_{AUC}$, (c) the experimental growth in combination of the two drugs, $Exp_{AUC}$ (d) and the theoretical additivity line, $Th_{AUC}$ (e) SIF. The five conditions passed the normality test.

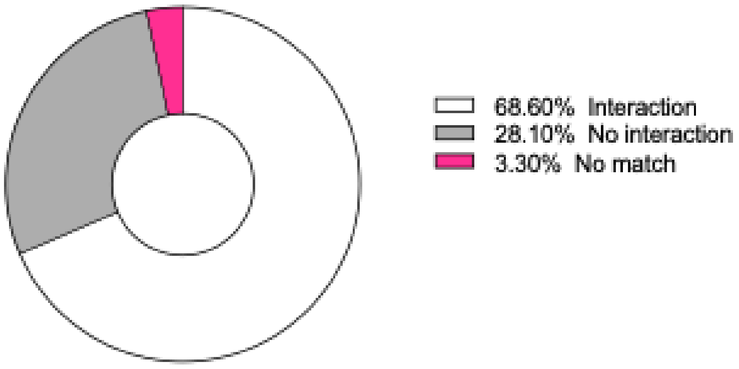


Supplementary figure S2: 121 isolate antibiotic combinations were assigned as interactions (either synergy or antagonism) or non-interactions (additive) according to the $Exp_{AUC}$ and $Th_{AUC}$ non-parametric t-test comparison and to SIF. Most of them were correspondingly classified as interactions (68.6%) or non-interactions (28.1%) with the two criteria, and only 4 of them (3.3 %) showed a mismatched classification.

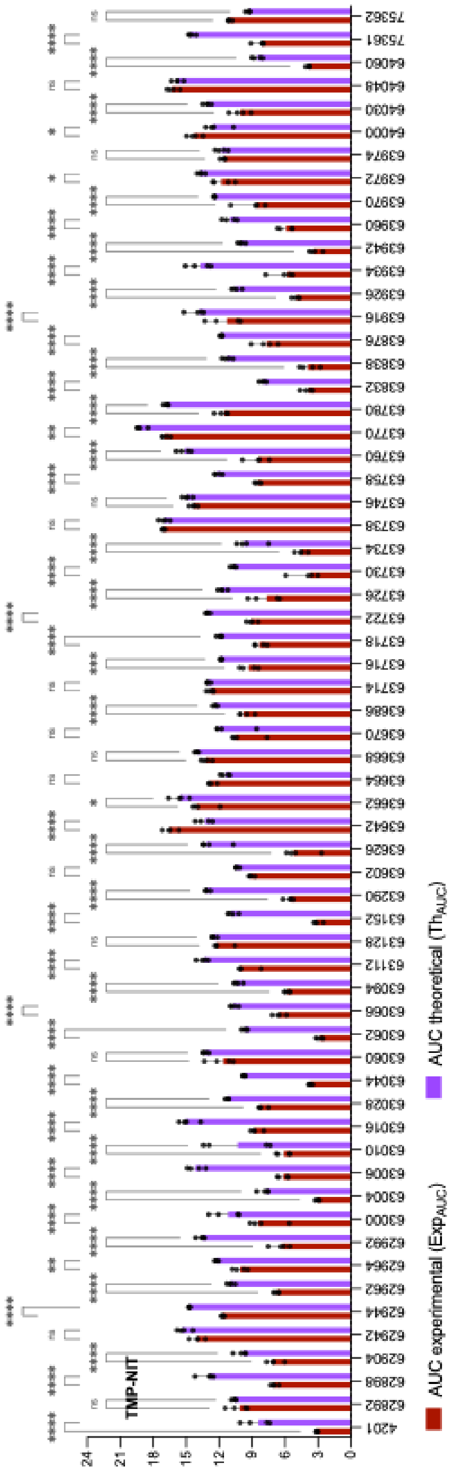


(a)

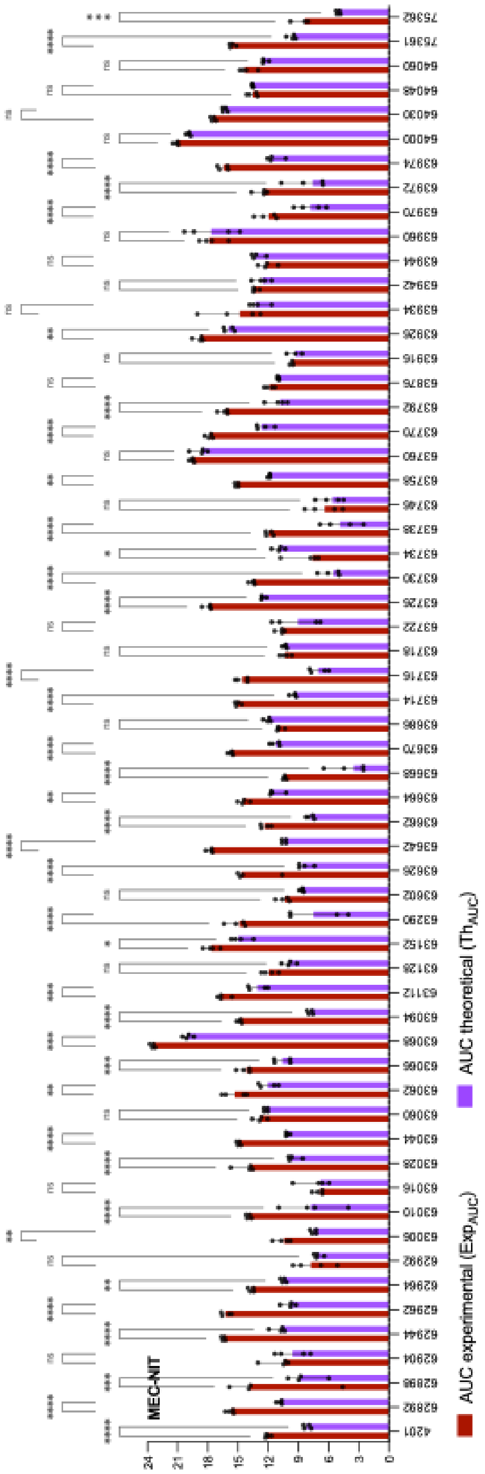


(b)

Supplementary figure S3: The AUC of the experimental relative line ($Exp_{AUC}$), in purple, and of the theoretical additivity line ($Th_{AUC}$), in dark red, are indicated for each isolate. The median values were determined from four biological replicates, marked with circles. Results obtained for the (a) TMP+NIT interaction, and (b) MEC+NIT interaction.

Significant differences between the AUC of Exp and Th lines were determined using nonparametric paired t-tests.

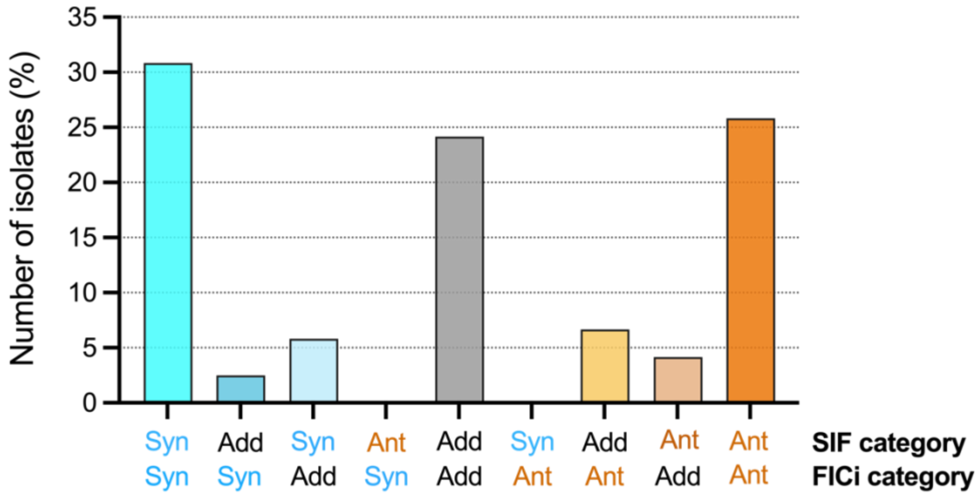


(a)

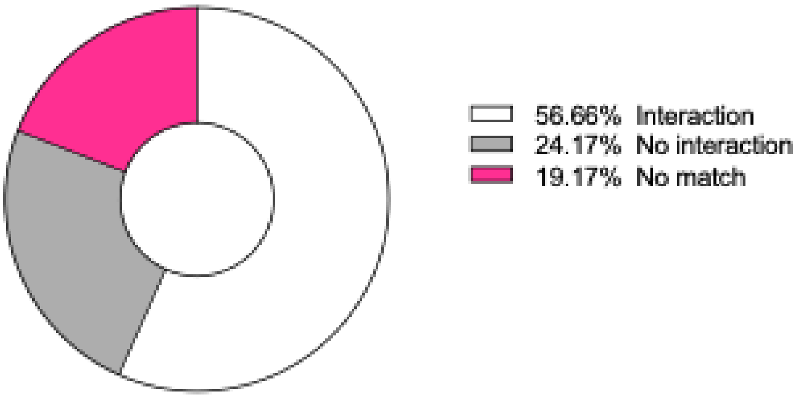


(b)

Supplementary figure S4: Comparison of the distribution by categories of the isolates according to SIF and FICi. (a) Overall, there is an agreement in the classification as synergistic (in blue), additive (in grey) or antagonistic (in orange) when each case is evaluated at lethal (using FICi) or sub-MIC (with SIF) conditions. (b) 80.83% isolates showed the same outcome with SIF and FICi (30,83% synergy, 24.17% additive and 25.83% antagonism). Among the 120 isolates tested, none of them showed an opposite phenotype when the two methodologies were compared.

| | | FICi | | |
|---|---|---|---|---|
| | | Syn | Add | Ant |
| SIF | Syn | 37 | 7 | 0 |
| | Add | 3 | 29 | 8 |
| | Ant | 0 | 5 | 31 |

$$Sensitivity = {}^{68}/_{80} \times 100 = 85.0\%$$

$$Specificity_{SYN} = {}^{73}/_{80} \times 100 = 91.25\ \%$$

$$Specificity_{ADD} = {}^{68}/_{79} \times 100 = 86.08\ \%$$

$$Specificity_{ANT} = {}^{76}/_{81} \times 100 = 93.83\ \%$$

$$Accuracy = {}^{97}/_{120} \times 100 = 80.83\ \%$$

Supplementary figure S5: Determination of the sensitivity and specificity of SIF, using FICi values as true classification. The sensitivity of SIF is 85.0 %, whereas the specificity was found to depend on the class, with an average of 90.6 %. The accuracy of SIF is 80.8 %.
